# Benchmarking Large Language Models for Transcriptomic Data Analysis and Visualization: Multi-Model Workflow Evaluation

**DOI:** 10.64898/2026.10.22.761023

**Authors:** Anjali Iyer, Arjun Subedi, Kimberly S. Butler, Raga Krishnakumar

## Abstract

This study presents a comprehensive benchmarking evaluation of large language models (LLMs) and agentic coding systems for automated analysis and visualization workflows for transcriptomic data, in particular human responses to COVID-19 infection. Multiple models, including ChatGPT 5.5, Claude Sonnet 4.6, Gemini 3.1, and Llama4-Scout, were assessed across a series of scientific prompts designed to test their ability to reproduce workflow components such as dimensionality reduction, clustering, outlier detection, reproducibility, and code generation. The benchmark protocol involved varying prompt specificity and examined both quantitative metrics such as clustering scores and outlier counts as well as qualitative aspects such as code quality and reproducibility. Results indicate that prompt specificity improves code structure and methodological correctness, but quantitative performance depends heavily on preprocessing choices and execution environment. While some models produced strong code and reproducible outputs, alignment with scientific targets like label-based clustering was inconsistent. The findings highlight the promise and limitations of current LLMs for automating scientific data analysis and visualization and suggest directions for future benchmarking and workflow standardization.

## 1 Introduction

### 1.1 Motivation

The arrival of big data in biology has fundamentally transformed the way hypotheses are tested and experiments are run. This transformation has accelerated the need for robust and scalable data analysis and visualization workflows in scientific research. As datasets grow in complexity and size, automating these workflows becomes increasingly important for timely and accurate insights. Large language models (LLMs) and agentic coding systems have emerged as powerful tools for generating scientific code and orchestrating data analysis pipelines, offering the potential to streamline tasks such as dimensionality reduction, clustering, outlier detection, and visualization.

Despite their exciting promise and potential, the scientific validity and reproducibility of LLM-generated workflows have not been established, and our ability to trust the outputs of the LLMs for high-consequence decision-making remains uncertain. In addition, each domain and each subset of tasks requires a distinct and careful evaluation and development of domain-specific metrics. Existing studies benchmarking the LLMs often focus on code generation or isolated tasks, but few systematically benchmark these models across the full spectrum of scientific workflow components (Cassano et al., 2023; Lai et al., 2023; Jimenez et al., 2024). Importantly, the effectiveness and reliability of LLMs in biological data analysis and data visualization are highly dependent on the prompts provided. This includes not only the stability of the final result, but the reproducibility of the methods which will allow rigorous evaluation and establish validity of LLM-generated scientific workflows.

While it is not possible to evaluate every possible prompt in every possible domain, studies are urgently required to shed light on how individual researchers might approach utilizing the AI in a responsible manner to achieve their goals. As LLM-prompting is increasingly becoming a primary means of conducting research, rigorous evaluation frameworks are needed. Because prompt performance is often highly task-dependent, effective evaluation must be grounded in the specific scientific use case rather than inferred from general benchmarking alone. This work uses a representative biological data science workflow to demonstrate one such evaluation framework for LLM-generated analyses that considers code quality, reproducibility, and alignment with scientifically meaningful performance metrics.

By designing a multi-model benchmark protocol with varying prompt specificity and comprehensive evaluation metrics, this study aims to identify strengths, limitations, and best practices for automated workflow generation for a common data science task as a proof-of-principle study. The ultimate goal is to inform future adoption of LLMs by scientists and researchers, ensuring that automation enhances, not compromises, scientific rigor and reproducibility.

### 1.2 Related Work

Biological datasets, especially molecular “omics” datasets, are often high-dimensional, noisy, heterogeneous, and sensitive to preprocessing choices. Standard analysis workflows commonly include normalization, dimensionality reduction, clustering, and visualization to identify structure in complex measurements such as gene expression, protein abundance, or clinical phenotypes. These steps are particularly central in single-cell and other high-capacity biological analyses, where methods such as principal component analysis (PCA), a linear dimensionality-reduction method that projects high-dimensional data onto a smaller number of orthogonal components capturing the largest sources of variance, and t-distributed stochastic neighbor embedding (t-SNE), a nonlinear visualization method that embeds high-dimensional samples into two or three dimensions while preserving local neighborhood structure, are routinely used to summarize variation and identify biologically meaningful subgroups (Luecken & Theis, 2019; Kiselev et al., 2019; Lähnemann et al., 2020). Because results can vary substantially with feature selection, scaling, parameter settings, batch effects, and software environment, reproducible workflow design is essential (Sandve et al., 2013; Deelman et al., 2009; Di Tommaso et al., 2017). In this study, COVID-19 data serve as a case study for evaluating the broader challenge of whether AI-generated biological data-analysis workflows are executable, scientifically valid, reproducible, and robust.

Large language models have recently become influential tools for software development and data-analysis assistance. Early work on code generation with language models showed that transformer-based systems can generate useful code from natural-language specifications, although their success depends on task formulation, sampling strategy, and problem complexity (Austin et al., 2021). More recent evaluations show that current frontier and specialized coding models now support increasingly seamless software-development assistance, such as code generation, revision, debugging, and realistic benchmark-based assessment (Nogueira, 2025; Kumarappan et al., 2026). However, performance remains sensitive to model size, benchmark design, task complexity, and workflow context (Hasan et al., 2026). In addition, recent work on collaborative coding systems suggests that as individual models become more capable, multi-agent collaboration may provide diminishing or task-dependent benefits (Kim et al., 2026).

However, domains are large and individual use-cases cannot all be explored. Therefore, it must be understood not only how to set up benchmarks but evaluate for what tasks models are variable, produce uncertainty and, make an individual (or collective) judgment on whether that is an acceptable amount of variability.

Several benchmarks have begun to evaluate code-generation systems under more realistic data-science and software-engineering conditions. (Cassano et al., 2023; Lai et al., 2023; Jimenez et al., 2024).

In addition, recent work has also shown growing interest in how agentic AI systems should be characterized, evaluated, and benchmarked, particularly as these systems move from single-response generation toward multi-step tool use and workflow execution. Studies of agentic AI workloads have begun to define the computational and behavioral characteristics of these systems, while broader surveys summarize emerging agent architectures, planning strategies, tool-use patterns, and evaluation challenges (Yuan et al., 2026; Plaat et al., 2026). However, despite this growing body of work, consensus has not yet emerged on best practices for benchmarking agentic scientific workflows.

Biological data analysis often requires chained operations rather than isolated code snippets. The emergence of science-focused AI tools such as Anthropic’s Claude Science(Anthropic, 2026) or GPT Rosalind (OpenAI, 2026) further reflects this broader movement from code assistance toward integrated research environments. While both systems place substantial emphasis on drug discovery, they are also relevant to broader questions about AI-assisted computational science because they seek to automate and streamline complex chains of research activity, and a detailed assessment of how the embedded language models perform will be critical to direct scientists on how to use them.

Reproducibility is central to scientific workflow automation because computational results must be not only efficient to generate, but also verifiable, repeatable, and methodologically sound. Foundational work in reproducible computational research argues that scientific claims should be supported by accessible data, executable code, and transparent documentation of the computational environment (Peng, 2011). Practical guidelines further emphasize version control, fixed random seeds, complete parameter reporting, and automated scripts that minimize manual intervention (Sandve et al., 2013).These concerns are especially relevant for LLM-generated workflows, where outputs may vary across prompts, model versions, decoding settings, and dependency configurations. For automated scientific workflows, this means that successful code execution alone is insufficient: evaluation must also determine whether generated workflows follow sound scientific and statistical practice, including appropriate handling of preprocessing, labels, scaling, and validation.

In this study, we not only augment the growing body of work evaluating LLMs for specific tasks, but also to provide a roadmap for researchers who may not be AI experts to feel enabled to design and execute their own benchmarking of LLMs before deciding on a path forward. This work focuses on a very small-scale task to highlight the need for benchmarking even when the task at hand is seemingly simple. In addition, this allows us to quantitatively and transparently evaluate the LLMs against a hard-coded gold-standard benchmark. This study builds on these strands of prior work by evaluating LLMs and agentic coding systems for automated transcriptomic data-analysis and visualization workflows. Unlike benchmarks focused only on short programming tasks or general-purpose reasoning, the proposed evaluation examines whether models can generate coherent end-to-end scientific workflows involving preprocessing, dimensionality reduction, clustering, outlier detection, visualization, and reproducibility documentation. The study’s emphasis on prompt specificity, execution environment, quantitative metrics, and qualitative code review reflects prior evidence that model outputs are sensitive to task formulation and that scientific validity cannot be assessed by executability alone. By situating LLM-generated analysis workflows within the broader literature on biological data infrastructure, code generation, agentic AI, reproducible computation, and benchmark design, this work addresses a gap in current evaluation practice and contributes toward more rigorous standards for AI-assisted scientific analysis.

## 2 Methods

### 2.1 Dataset Description

The benchmark utilized a comprehensive COVID gene-expression dataset compiled from multiple studies. Every study has publicly available data with known accession numbers. Below is a table of all four studies and their corresponding accession number (Wang et al., 2023; Overmyer et al., 2020; Thompson et al., 2023; McClain et al., 2021). For simplicity, the study names used here are the four last names of the first author for the original publications.

The goal of utilizing these four studies is to emulate a pipeline that is likely to be carried out by a researcher asking whether the effects of COVID19 on gene expression is strong enough to overcome the study-specific batch effect. We chose these studies because we know the outcome of this analysis, and we can evaluate the outputs of the LLMs accordingly. The dataset is transposed such that genes are represented as rows and samples as columns, with each sample labeled as either COVID or non-COVID. Most benchmarking runs treated the data as comprising 1,895 COVID samples and 209 non-COVID samples.

Prior to model evaluation, a hard-coded pipeline was executed on the dataset to generate “Gold Standard” results. This pipeline included PCA dimensionality reduction, PCA + t-SNE visualization, KMeans clustering, Local Outlier Factor (LOF) outlier detection. K-means clustering is an unsupervised learning method that partitions observations into k clusters by assigning each sample to the cluster with the nearest centroid, thereby minimizing within-cluster variation (Jain, 2010). LOF is an unsupervised anomaly-detection method that identifies samples with substantially lower local density than their nearest neighbors, flagging those samples as potential outliers (Breunig et al., 2000). These results served as the baseline for all subsequent model comparisons. The code for this pipeline is provided with this paper.

### 2.2 Model Selection

The benchmark evaluated four large language models, each chosen for their publicly documented strong performance on code-generation tasks and their representation of different contemporary model families. The four models were ChatGPT 5.5, Claude Sonnet 4.6, Gemini 3.1, and Llama4-Scout. These models were selected based on their prominence in public model releases and technical capabilities available at the time of study design. Together, they span both proprietary and open-weight systems and reflect variation in architecture, deployment model, and reported coding performance. This diversity enabled a robust comparison of methodological correctness, code quality, and reproducibility across a representative set of state-of-the-art LLM systems.

For this study, each of these large language models were used as direct prompt-response systems. Following each prompt, each LLM was prompted to generate a complete analysis script, which was then executed manually to produce the resulting plots and text outputs. Thus, the LLM-generated code functioned as an intermediate artifact between the prompt and the final analysis results. The primary objective of using four different LLMs was not to quantify variability across repeated generations of code from the same prompt, but to evaluate whether each model, under different prompt specificity conditions could generate executable workflows whose outputs aligned with those of the hard-coded benchmark pipeline. Code quality and reproducibility were evaluated as supporting dimensions of workflow usability, while the main source of variation examined in this study was differences in outputs across models and prompt versions. More details on prompt specificity and evaluation metrics will be included in following sections.

All computational workflows in this study were implemented in Python, using a standard data-science stack centered on pandas for data handling, scikit-learn for dimensionality reduction, clustering, and outlier detection, and matplotlib for visualization. The LLMs were prompted to generate Python scripts compatible with this environment so that outputs could be compared consistently against the hard-coded benchmark pipeline.

### 2.3 Benchmark Design and Rationale

The benchmark was designed to systematically evaluate the capabilities of large language models (LLMs) and agentic coding systems in automating COVID-19 data analysis and visualization workflows. The overarching goal was to move beyond superficial code generation metrics and instead probe both the scientific validity and practical reproducibility of LLM-driven workflows in a real-world context.

### 2.4 Prompt Coverage

Seven scientific prompts were developed to systematically cover the core components of the data-analysis pipeline. These prompts evaluated dimensionality reduction, clustering, outlier detection, reproducibility, and code-generation quality.

The seven prompts were defined as follows:

**Prompt 1:** PCA visualization
**Prompt 2:** PCA + t-SNE visualization
**Prompt 3:** K-means clustering
**Prompt 4:** Full Test— PCA + t-SNE + K-means
**Prompt 5:** LOF outlier detection
**Prompt 6:** Reproducibility test

**Prompt 7:** Code-generation quality Taken together, these prompts assessed the major analysis components of interest:

**Dimensionality Reduction:** Prompts 1, 2, and 4
**Clustering:** Prompts 3 and 4 **Outlier Detection:** Prompt 5 **Reproducibility:** Prompt 6
**Code Generation Quality:** Prompt 7

Running through all seven prompts were the experiments that tested all parts of the hard-coded pipeline.

To probe the impact of instruction specificity, each prompt was tested in three versions: general, informed, and expert. Each version of the prompt was developed and evaluated manually before being employed for testing. For the general prompt, minimal instruction was provided with the aim of testing the model’s default reasoning and code synthesis. For the informed prompt, dataset orientation was provided along with label information, and context to guide the model toward correct data handling. The expert prompt made the workflow expectations fully explicit by specifying file-loading procedures, preprocessing steps, random-seed settings, parameter choices, and expected output formats, which aimed to mirror the level of detail found in published scientific protocols. In practical terms, the expert prompt was designed to state explicitly every step that one might otherwise assume the LLM already knew to perform. This framing allowed prompt specificity to be evaluated systematically, from under-specified requests to highly explicit workflow instructions.

An example of the 3 varying levels of specificity of prompts for PCA visualization (prompt 1) is as follows:

### v1 - General

I have a COVID gene expression dataset at /path/to/file. Visualize it using PCA. Activate /project/venv first. Save to /path/to/output.

### v2 - Informed

I have a COVID gene expression dataset at /path/to/input/file. The file is transposed — genes are rows, samples are columns. Label column is class _label (COVID or non-COVID). Run PCA and plot the top 2 components colored by labels. Activate /project/venv first. Save to /path/to/output.

### v3 - Expert

I have a COVID gene expression dataset at path/to/file. Load with index_col = 0, transpose so samples are rows, extract class_label (COVID or non-COVID) as labels, drop from features, convert to numeric filling NaN with 0. Run PCA with 50 components random_state=42. Plot PC1 vs PC2: COVID=red (#FF0000), non-COVID=blue (#0000FF), with legend, axis labels, and title. Activate /project/venv first. Save to /path/to/output.

### 2.5 Scientific Rigor

This multi-version approach allowed for assessment of both methodological correctness and adaptability to scientific requirements. The rationale was to capture not only quantitative performance but also qualitative aspects such as code structure, reproducibility, and scientific alignment.

By evaluating models across these axes, the benchmark provides actionable insights for both LLM developers and all other end-users.

### 2.6 Workflow Overview

The entire benchmarking process included four overarching general steps: Generate pipeline gold standard once before proceeding with any other step, run each version of every prompt across all LLMs, fill out comparison table, and log each run. The primary dataset used in this study, contained gene expression profiles from multiple studies with each sample labeled as ‘COVID’ or ‘non-COVID’.

### Step 1: Generating the pipeline "Gold Standard" results

Before testing any LLM, the hard-coded pipeline was run on the COVID dataset to generate the results that all other model results would be benchmarked against. This was done by running all the baseline experiments: PCA plotting, PCA + t-SNE visualization, KMeans clustering, PCA + t-SNE + KMeans visualization, and LOF outlier detection, on the COVID-19 dataset using the hard-coded pipeline and using all generated results from these baseline experiments to generate what was known as the “Gold Standard” results. LLM-generated results were benchmarked against these “Gold Standard” results to evaluate their performance. It is important to note that Prompts 6 and 7 were not explicit components of the hardcoded pipeline and therefore were not included in the “gold-standard” results. Instead, these prompts were designed to assess reproducibility and code-generation quality as standalone model capabilities, providing a supplementary evaluation alongside the primary comparisons against the gold-standard pipeline.

### Step 2: Model testing on every prompt

The workflow then proceeded through the series of prompts and versions as described previously with the results from every run being saved for future evaluation of visual and quantitative quality. All seven prompts were run for all models tested. Prompts 1-5 generated results either in the form of visual plots, scores, or both, that would later be compared against the “Gold Standard” results.

As mentioned previously, four models were included in the benchmark: ChatGPT 5.5, Claude Sonnet 4.6, Gemini 3.1, and Llama4-Scout. Each model was evaluated across seven prompts, and completion status was tracked for every workflow component. All four models successfully completed Prompts 1–7. This design enabled a structured comparison across models and prompt conditions; strengths and limitations were then inferred from later evaluation of the generated workflows, including execution success, code quality, reproducibility, and agreement with the gold-standard pipeline.

### Step 3: Fill out comparison table

For all prompts with quantitative results which are prompts 3 – 7, for every model, a comprehensive comparison table was completed to make analysis on model performance across each prompt clearer and easier.

The quantitative metrics used in evaluation were LOF outlier count, Silhouette Score, ARI, Code quality score, and a binary “yes” or “no” response to the reproducibility test matching question, essentially, did each run result in identical results. reproducibility score (“yes” or “no”). A table outlining each of these metrics along with why each metric was used is as described in Table 3.

**Table 1:** NCBI GEO Accession Number

| Study name | Accession number |
| --- | --- |
| Wang | GSE217948 |
| Overmyer | GSE157103 |
| Thompson | GSE215865 |
| McClain | GSE161731 |

**Table 2:** Comparison table template

| Prompt | Metric | Pipeline score | Prompt v1 | Prompt v2 | Prompt v3 |
| --- | --- | --- | --- | --- | --- |
| 3 KMeans | Silhouette |  |  |  |  |
| 3 KMeans | ARI |  |  |  |  |
| 4 Full | Silhouette |  |  |  |  |
| 4 Full | ARI |  |  |  |  |
| 5 LOF | Outliers |  |  |  |  |
| 6 Repro | Match? | N/A |  |  |  |
| 7 Code | Quality /25 | N/A |  |  |  |

**Table 3:** Evaluation categories and metrics used to assess LLM-generated workflows.

| Evaluation category | Metric | Interpretation |
| --- | --- | --- |
| Clustering | Silhouette Score | Higher means more separated clusters |
| Clustering | Adjusted Rand Index (ARI) | Agreement with true COVID/non-COVID labels |
| Outlier detection | LOF outlier count | Compared to pipeline count |
| Reproducibility | Yes/No match | Repeated runs produced same outputs |
| Code quality | 25-point rubric | Input checks, parameters, docs, seeds, correctness |

The empty comparison table followed the template outlined in Table 2.

### Step 4: Log every run

Finally, for each model, every version of every run was marked a “pass” or “fail” with notes describing why that designation was made for further analyses purposes. The organization of this table consisted of the following column headers: “Prompt #”, “Model”, “Version”, “Pass/Fail”, “Attempts, “Time”, “Score”, and “Notes”. “Score” held any associated numerical scores of that particular prompt and version (silhouette score, ARI, LOF, rubric score).

## 3 Results

### 3.1 Evaluation Metrics

To comprehensively assess performance across all models, both quantitative and qualitative evaluation metrics were used. To qualitatively assess PCA and t-SNE plotting results, a visual checklist was employed where it was checked that the plots contained the correct colors, a legend, labeled axes, and correctly separated clusters.

Quantitative results were used to assess model performance for clustering and outlier detection. Clustering metrics included silhouette score which measures intra-cluster cohesion and inter-cluster separation where higher values indicate better-defined clusters and adjusted rand index (ARI) which quantifies the agreement between predicted clusters and true COVID/non-COVID labels where ARI = 1.0 is perfect, 0 is random, and negative values indicate anti-alignment. It is important to note here that models were not ranked based on whether Silhouette Score alone increased or decreased. Instead, performance was interpreted relative to the benchmark pipeline using the paired behavior of Silhouette Score and ARI. Higher Silhouette with low ARI indicated well-separated but biologically misaligned clusters, whereas a somewhat lower Silhouette with ARI closer to the pipeline indicated outputs that better matched the intended COVID/non-COVID structure. Thus, rankings reflected closeness to the benchmark profile, not simply whether individual metric values were numerically larger.

The outlier detection metric included LOF outlier count which measured the number of samples that were flagged as outliers by Local Outlier Factor which was then compared to the outlier count calculated by our gold-standard pipeline.

Code quality was measured using a 25-point rubric manually developed for this specific study where scripts were assessed on whether they ran without error, exposed configurable parameters, performed basic input validation or precondition checks, included explanatory comments, and set random seeds where relevant. For each rubric category, scripts were scored from 0 to 5, where 0 indicated that the category was not addressed, 3 indicated that the element was present but applied inconsistently, and 5 indicated that the element was implemented consistently and contributed meaningfully to the script’s functionality The codequality evaluation was intentionally scoped to practical properties relevant to small scientific analysis scripts, including executability, configurability, transparency, and reproducibility. It did not assess broader software-engineering attributes such as modularity or maintainability.

To complement the rubric-based assessment of code-generation quality, after testing all models and recording the corresponding results, a static analysis was also performed using Ruff, a Python linter and code-quality tool that includes support for checking McCabe cyclomatic complexity. Cyclomatic complexity is a widely used software metric that estimates the structural complexity of a function by counting the number of linearly independent execution paths through the code; higher values generally indicate code that may be more difficult to inspect, test, or maintain (McCabe, 1976). In this study, we used a maximum complexity threshold of 6 as a practical cutoff for small LLM-generated scientific scripts, consistent with the goal of favoring code that remained straightforward to interpret and reuse. This metric was used only as a supplementary indicator of structural complexity and not as a standalone measure of scientific or software quality.

Reproducibility was measured by evaluating with a binary ‘yes’ or ‘no’ if repeated runs with fixed seeds produced identical results.

### 3.2 Individual Model Results

**Pipeline Gold Standard Results** The reference pipeline established baseline metrics for all subsequent model comparisons:

- Clustering:

**–** K-means Silhouette Score: 0.513
**–** Full test Silhouette Score: 0.5582
**–** K-means ARI: 0.410
**–** Full test ARI: 0.413
- Outlier Detection:

**–** LOF outlier count: 58

### ChatGPT 5.5 Results

ChatGPT 5.5 successfully completed Prompts 1-7. As seen in Table 4, Silhouette scores were notably higher than pipeline scores going up to 0.858, but ARI values were often negative or near zero, indicating that while models may have successfully identified some sort of structure, in this case clusters, the identified clusters did not align with COVID/non-COVID labels. It performed well at outlier detection with a LOF outlier count of 56, which was very close to the pipeline’s count of 58.

**Table 4:** Quantitative benchmark results

| Prompt | Metric | Pipeline | Prompt v1 | Prompt v2 | Prompt v3 |
| --- | --- | --- | --- | --- | --- |
| 3 K-means | Silhouette | 0.513 | 0.858 | 0.858 | 0.410 |
| 3 K-means | ARI | 0.410 | -0.00698 | -0.00698 | 0.0514 |
| 4 Full | Silhouette | 0.558 | 0.858 | 0.452 | 0.410 |
| 4 Full | ARI | 0.413 | -0.00698 | 0.193 | 0.051 |
| 5 LOF | Outliers | 58 | 94 | 211 | 56 |
| 6 Reproducibility | Match? | N/A | Yes | Yes | Yes |
| 7 Code generation | Quality /25 | N/A | 22 | 25 | 18 |

The results displayed in Table 4 were interpreted relative to the hard-coded pipeline baseline rather than by assuming that larger Silhouette Score and ARI values alone always proved better performance in these categories. For the clustering and full test tasks, performance was assessed by how closely the paired

Silhouette Score and ARI values matched the benchmark pipeline. In Prompt 3, Versions 1 and 2 increased the Silhouette Score by +0.345 relative to the pipeline (0.858 vs 0.513) but decreased the ARI by *−*0.417 (*−*0.00698 vs. 0.410), which indicated well-separated but biologically misaligned clusters. Version 3, in contrast, decreased Silhouette Score by –0.103, (0.410 vs 0.513) but reduced the ARI gap to –0.359 (0.0514 vs 0.410, which made it a closer overall match to the benchmark profile. Prompt 4 showed a similar pattern, but there was a slight dip in performance from Version 2 to Version 3, both of these versions however, still outperformed Version 1, where Version 2 reduced the Silhouette difference to –0.106 and the ARI difference to –0.220, which yielded the closest overall match to the pipeline. In contrast, as can be seen in Table 5, code-generation quality declined from Version 2 to Version 3, suggesting that increased prompt specificity improved alignment with benchmark analysis outcomes in some tasks, but not uniformly across all evaluation dimensions.

**Table 5:** Code-generation quality rubric scores across prompt versions.

| Criteria | v1 | v2 | v3 |
| --- | --- | --- | --- |
| Runs without error | 5 | 5 | 0 |
| Parameters configurable | 5 | 5 | 5 |
| Input validation checks present | 5 | 5 | 3 |
| Has docstrings/comments | 5 | 5 | 5 |
| Uses random seeds | 3 | 5 | 5 |
| Total /25 | 22 | 25 | 18 |

Apart from the quantitative observations, PCA/t-SNE plots showed good visual separation, but axis scaling sometimes differed from the pipeline and code structure improved markedly with expert-level prompts.

### Claude Sonnet 4.6 Results

Claude Sonnet 4.6 also successfully completed Prompts 1–7. Silhouette and ARI scores were consistently different from the recorded pipeline scores. Outlier count was recorded slightly inconsistently across varying levels of prompt specificity, but all recorded more outliers than the pipeline had originally recorded.

Based on the Silhouette Score and ARI score pairing interpretation described for the previous model, it can be seen in Table 7 that Claude had very little change in performance over prompts 3,4, 5, and 7 across prompt versions, but achieved slight improvement in outlier detection count from prompt version 1 to version 3.

**Table 6:** Reproducibility results across prompt versions.

| Version | Run 1 Silhouette | Run 2 Silhouette | Match? |
| --- | --- | --- | --- |
| v1 | 0.858 | 0.858 | Yes |
| v2 | 0.858 | 0.858 | Yes |
| v3 | 0.410 | 0.410 | Yes |

**Table 7:** Quantitative benchmark results for Claude Sonnet 4.6 across prompt versions.

| Prompt | Metric | Pipeline | Prompt v1 | Prompt v2 | Prompt v3 |
| --- | --- | --- | --- | --- | --- |
| 3 K-means | Silhouette | 0.5130 | 0.8600 | 0.8600 | 0.8600 |
| 3 K-means | ARI | 0.4097 | -0.0070 | -0.0070 | -0.0070 |
| 4 Full | Silhouette | 0.5582 | 0.8600 | 0.8600 | 0.8600 |
| 4 Full | ARI | 0.4129 | -0.0070 | -0.0070 | -0.0070 |
| 5 LOF | Outliers | 58 | 106 | 106 | 94 |
| 6 Reproducibility | Match? | N/A | Yes | Yes | Yes |
| 7 Code generation | Quality /25 | N/A | 25 | 22 | 25 |

The model also produced reproducible scripts across runs for all prompts as can be seen in Table 8.

**Table 8:** Reproducibility results for Claude Sonnet 4.6 across prompt versions.

| Version | Run 1 Silhouette | Run 2 Silhouette | Match? |
| --- | --- | --- | --- |
| v1 | 0.860 | 0.860 | Yes |
| v2 | 0.860 | 0.860 | Yes |
| v3 | 0.410 | 0.410 | Yes |

Once again, while code generation was strong overall as can be seen in Table 9, clustering in PCA and t-SNE plots was very different than those produced by the pipeline.

**Table 9:**
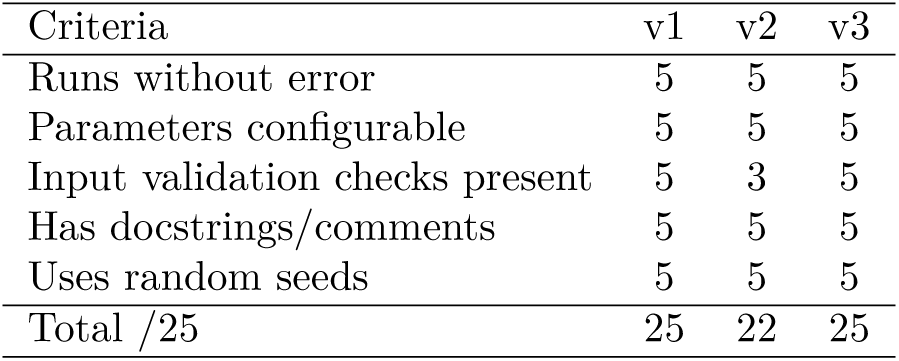
Code-generation quality rubric scores for Claude Sonnet 4.6 across prompt versions.

| Criteria | v1 | v2 | v3 |
| --- | --- | --- | --- |
| Runs without error | 5 | 5 | 5 |
| Parameters configurable | 5 | 5 | 5 |
| Input validation checks present | 5 | 3 | 5 |
| Has docstrings/comments | 5 | 5 | 5 |
| Uses random seeds | 5 | 5 | 5 |
| Total /25 | 25 | 22 | 25 |

### Gemini 3.1 Results

Gemini 3.1, like all previous models completed Prompts 1–7.

As can be seen in Table 10, KMeans clustering scores were inconsistent, with errors resulting in the first version prompt that prevented scores to be generated, but the expert level version prompt produced both a Silhouette Score and ARI that were very similar to those of the pipeline. Clusters looked to be well separated from the plots but did not align with true labels. The LOF outlier count was consistently generated to be 56, similar to the count of 58 recorded by the pipeline. Gemini 3.1 had among the best handling of quantitative tasks and once again highlighted how increased specificity can improve results particularly in reference to the KMeans experiment ARI and Silhouette scores from versions 1 to versions 3.

**Table 10:** Quantitative benchmark results for Gemini 3.1 across prompt versions.

| Prompt | Metric | Pipeline | Prompt v1 | Prompt v2 | Prompt v3 |
| --- | --- | --- | --- | --- | --- |
| 3 K-means | Silhouette | 0.5130 | N/A | 0.8584 | 0.4096 |
| 3 K-means | ARI | 0.4097 | -0.0070 | -0.0070 | 0.0514 |
| 4 Full | Silhouette | 0.5582 | 0.4096 | 0.4096 | 0.4096 |
| 4 Full | ARI | 0.4129 | 0.0514 | 0.0514 | 0.0514 |
| 5 LOF | Outliers | 58 | 56 | 56 | 56 |
| 6 Reproducibility | Match? | N/A | Yes | Yes | Yes |
| 7 Code generation | Quality /25 | N/A | 23 | 25 | 20 |

In Table 11, It is displayed that the code scored a 25/25 on the rubric for v2 prompts; code was reusable, well-parameterized, and included explanatory comments. However, like with ChatGPT 5.5, code quality appeared to decline from version 2 to version 3, once again highlighting a potential tradeoff between improved quantitative task completion and “clean” code.

**Table 11:** Code-generation quality rubric scores for Gemini 3.1 across prompt versions.

| Criteria | v1 | v2 | v3 |
| --- | --- | --- | --- |
| Runs without error | 5 | 5 | 0 |
| Parameters configurable | 5 | 5 | 5 |
| Input validation checks present | 5 | 5 | 5 |
| Has docstrings/comments | 5 | 5 | 5 |
| Uses random seeds | 3 | 5 | 5 |
| Total /25 | 23 | 25 | 20 |

All runs with fixed seeds were fully reproducible as can be seen in Table 12.

**Table 12:** Reproducibility results for Gemini 3.1 across prompt versions.

| Version | Run 1 Silhouette | Run 2 Silhouette | Match? |
| --- | --- | --- | --- |
| v1 | 0.410 | 0.410 | Yes |
| v2 | 0.410 | 0.410 | Yes |
| v3 | 0.410 | 0.410 | Yes |

While visualizations were clear and well-labeled, cluster separation and axis scaling was very different from the pipeline’s clustering plots, however, code structure was robust across prompt versions.

### Llama4-Scout Results

As can be seen in Table 13, silhouette Score went all the way up to 0.8584 and ARI was consistently generated to be –0.0070 which indicated that clusters did not correspond to COVID/non-COVID labels. LOF outlier count was either generated to be 94 or 106 based on the prompt version, meaning the model consistently overestimated the number of outliers when compared to the pipeline.

**Table 13:** Quantitative benchmark results for Llama4-Scout across prompt versions.

| Prompt | Metric | Pipeline | Prompt v1 | Prompt v2 | Prompt v3 |
| --- | --- | --- | --- | --- | --- |
| 3 K-means | Silhouette | 0.5130 | 0.8584 | 0.8584 | 0.8584 |
| 3 K-means | ARI | 0.4097 | -0.0070 | -0.0070 | -0.0070 |
| 4 Full | Silhouette | 0.5582 | 0.8584 | 0.8584 | 0.8584 |
| 4 Full | ARI | 0.4129 | -0.0070 | -0.0070 | -0.0070 |
| 5 LOF | Outliers | 58 | 94 | 106 | 94 |
| 6 Reproducibility | Match? | N/A | Yes | Yes | Yes |
| 7 Code generation | Quality /25 | N/A | 17 | 17 | 23 |

It is seen in Table 14 that scripts produced by this model scored 17–23/25, so code was generally correct but sometimes lacked basic validation checks or parameterization. Notably, code quality score increased across prompts, but scores for the quantitative tasks did not increase or changed slightly, still remaining far off from the hard-coded pipeline’s results.

**Table 14:** Code-generation quality rubric scores for Llama4-Scout across prompt versions.

| Criteria | v1 | v2 | v3 |
| --- | --- | --- | --- |
| Runs without error | 5 | 5 | 3 |
| Parameters configurable | 2 | 2 | 5 |
| Input validation checks present | 0 | 0 | 5 |
| Has docstrings/comments | 5 | 5 | 5 |
| Uses random seeds | 5 | 5 | 5 |
| Total /25 | 17 | 17 | 23 |

While plots showed strong separation, cluster labeling was inconsistent with ground truth.

## 4 Conclusions

A consistent overall trend was that increasing prompt specificity generally improved model performance. General prompts produced the weakest results, informed prompts were more reliable, and expert prompts most often yielded the strongest performance, particularly on quantitative evaluation metrics.

However, this improvement was not uniform across all categories. One notable exception was code-generation quality, assessed using a 25-point subjective rubric evaluating whether scripts ran without error, allowed parameter configuration, included basic input validation checks, used docstrings or comments, and set random seeds. For several models, code quality declined from Version 2 to Version 3, indicating that although expert prompts often improved execution and task completion, they did not always produce cleaner or better-structured code. Among the models tested, Claude Sonnet 4.6 and Gemini 3.1 achieved the highest code-quality scores, each receiving 25/25 for generating robust, reusable, and well-documented scripts.

As briefly touched on, previously to supplement the rubric-based evaluation of code-generation quality, a static analysis of the generated Python scripts were performed using Ruff, a Python linting and code-quality tool. Higher complexity values generally indicate code that may be more difficult to inspect, test, or maintain. Because the generated workflows in this study were relatively small analysis scripts, we used a maximum complexity threshold of 6 as a conservative heuristic for identifying scripts whose control flow had become less straightforward. Most scripts fell below this threshold. Only four exceeded it: three Gemini 3.1 LOF scripts, each with a main function complexity of 7, and one Llama4-Scout script from the code-generation-quality task, in which the load_and_prepare function had a complexity of 10. These results suggest that most generated scripts were structurally simple by this metric, while a small subset exhibited more branching logic than desired for compact, easily inspectable workflows.

Of the quantitative tasks, the most challenging proved to be K-means clustering, the full test, and LOF outlier detection. Even when generated scripts executed successfully, their outputs were often only partial or weak by quantitative criteria. Despite this, ChatGPT 5.5 and Gemini 3.1 produced LOF outlier counts of 56, closely matching the pipeline baseline of 59, and therefore showed the strongest quantitative alignment for anomaly detection among the models evaluated.

For clustering tasks, all models were able to generate clusters with relatively high Silhouette Scores, indicating internally well-separated clusters. However, Adjusted Rand Index (ARI) values were often near zero or negative, showing that these clusters did not align with the true COVID/non-COVID labels. This distinction is important: the Silhouette Score measures geometric cluster separation, whereas ARI measures agreement with the biologically meaningful labels of interest. As a result, high Silhouette Scores did not necessarily indicate scientific success. In many cases, the models identified the clusters in the data, but not the relevant clusters (COVID vs. Non-COVID) that the task was intended to recover.

Visual inspection of the PCA and t-SNE outputs reinforced this conclusion. Although most models produced plots with apparent sample separation, only the hard-coded pipeline consistently generated groupings that aligned with the true biological labels. Taken together, these findings show that visual separation and successful execution alone are insufficient measures of scientific validity, and they underscore the importance of evaluating LLM-generated workflows against quantitative benchmarks.

## 5 Discussion

The benchmarking results reveal that LLMs and agentic coding systems can generate executable code and reproducible outputs for COVID data analysis and visualization workflows. However, scientific validity depends on more than code execution. Prompt specificity enhances code structure and explicitness but does not guarantee quantitative alignment with scientific targets. High Silhouette Scores alone are insufficient; ARI is essential for evaluating agreement with COVID/non-COVID labels. The strongest quantitative matches were observed in LOF outlier detection, suggesting that anomaly detection tasks may be more tractable for current models.

Generally, more detailed prompts generally helped improve scores, but there were tradeoffs, particularly for more complex prompts such as the full test and the qualitatively evaluated prompts, which where the PCA visualization and PCA + tSNE visualization prompts. Better task completion did not always mean better code quality, and executable code still did not consistently translate to results that were scientifically valid when compared against our benchmarking results. Furthermore, strong performance on this code-quality rubric should not be interpreted as evidence that LLM-generated code satisfies the broader requirements of production-quality scientific software. And although later testing of cyclomatic complexity provided an objective measure of structural complexity, it should be interpreted cautiously: low-complexity code can still be scientifically incorrect, and more complex code may sometimes reflect legitimate preprocessing or validation logic.

Code quality and reproducibility are promising, but limitations in clustering and batch-awareness highlight areas for improvement. Execution environment constraints, such as timeouts, can significantly impact benchmarking outcomes and should be considered in future evaluations.

### 5.1 Limitations and Open Questions

While comprehensive, there still exist several limitations that may affect interpretation of this benchmark. These most notably include the fact an eighth batch awareness prompt was not completed or included for most models as originally planned, the fact that visual comparisons of the PCA plotting and t-SNE + PCA plotting are subjective and qualitative, and the fact that only specific versions of some of these LLMs were available and that future updates of these models may yield different results. It is also important to factor in the possibility that preprocessing differences rather than model quality may influence quantitative results.

In addition to these limitations, some of the most notable questions that remain open to be answered after this study include:

- How can prompt design be further optimized for scientific alignment?
- What preprocessing standards should be adopted for benchmarking?
- How can execution environment constraints be mitigated?

## 6 Future Outlook

Best practices learned from the study included continuous benchmarking against a gold-standard pipeline, specifically, always comparing LLM-generated workflows to a rigorously validated reference to ensure scientific validity. AI should be used as a workflow accelerator rather than as a replacement for scientific validation. Future workflows should begin with a reliable baseline such as a known-good script or prior validated pipeline, so that LLM-generated outputs can be compared against an established reference. Prompts should be made as explicit as possible, specifying file structure, labels, preprocessing steps, expected outputs, and parameter settings, rather than assuming the model will infer these details correctly.

Reproducibility should also be enforced deliberately. Generated workflows should set random seeds, save metrics and plots, and record environment details to support repeatable execution and potential adaption to other similar analyses workflows. At the same time, outputs must be validated scientifically rather than judged only by whether code runs or plots appear reasonable. This includes always evaluating results against domain-relevant quantitative metrics, in this study those metrics included Silhouette Score, ARI, and LOF outlier detection, and confirming that generated analyses align with the underlying biological question. Because executable code can still embed flawed assumptions, generated scripts should also be inspected directly for preprocessing choices, label handling, parameter use, and other analysis decisions. Finally, failures should be documented, since unsuccessful runs can reveal hidden model assumptions, workflow fragility, and environment-specific limitations.

The development of biology-specific agents is aiming to create frameworks that are inherently previously benchmarked for specific analysis workflows. Examples are rapidly growing, and as these tools become more broadly available for use, benchmarking standards become increasingly important. While these tools are internally benchmarked, their wide adoption means that scientists will have to empower themselves to a) understand where likely sources of variability, uncertainty and fault are, and b) how to evaluate the tools for themselves to decide which one best suits their needs. We therefore suggest using benchmarks such as the ones we present here to test a range of agents and languages prior to making any biological determinations, future work should also focus on multi-study evaluations, developing strategies to handle environment-specific limitations and ensure portability of LLM-generated code, and incorporating additional models such as Devstral or future releases of models already included for expanded model coverage.

Overall, the adoption of LLMs in scientific research should be guided by comprehensive benchmarking, transparent reporting, and a commitment to continuous re-evaluation. Used carefully, these systems can accelerate analysis, but only by rigorously benchmarking both the capabilities and limitations of these systems, it can be ensured that these systems enhance rather than compromise scientific rigor and reproducibility.

**Figure 1:**
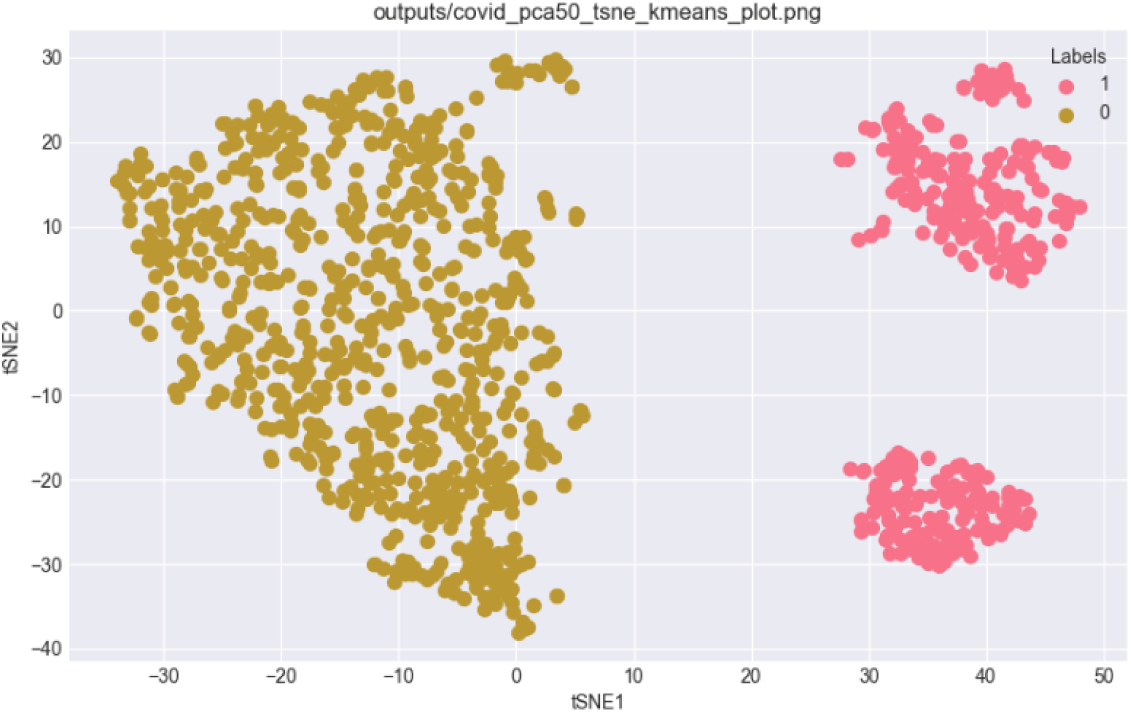
Pipeline gold-standard result for the Full Test experiment.

**Figure 2:**
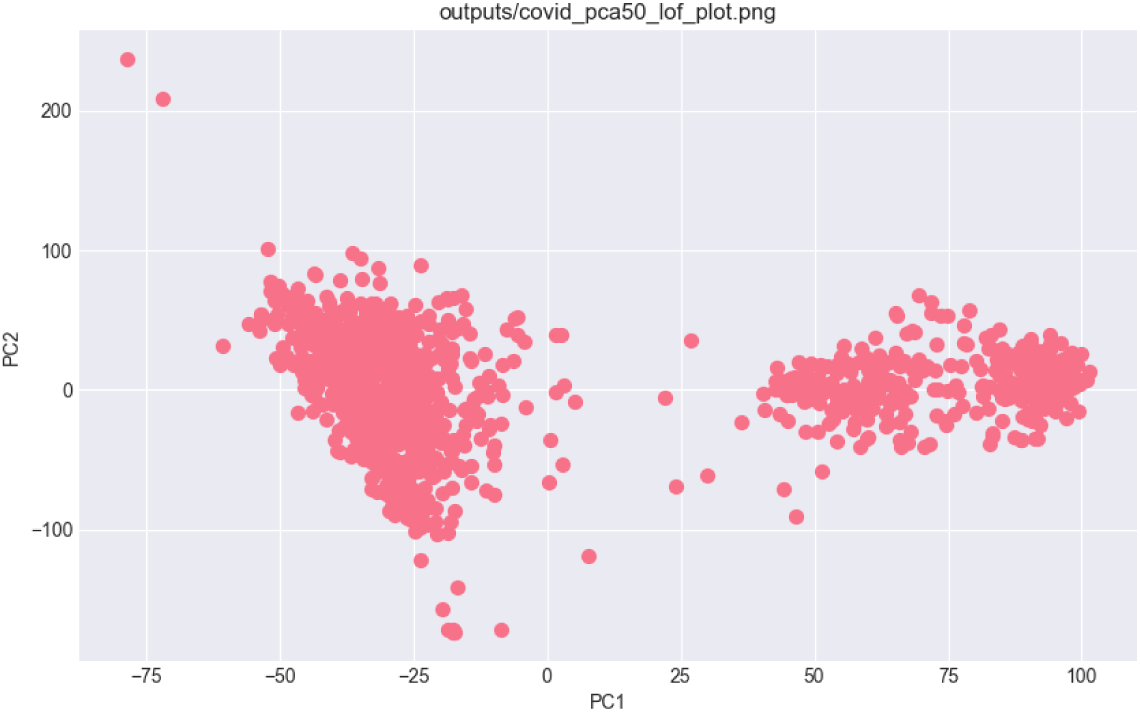
Pipeline gold-standard result for the LOF experiment.

## Acknowledgments

We thank Mark Forrer and Laura Weinstock for their comments on this manuscript. We also thank Jerilyn A. Timlin, Yooli K. Light and Patrick D. Finley for technical feedback during the project. This work was supported by the DOE Bioenergy Technology Office (BETO) and the Laboratory Directed Research and Development program (project 233091) at Sandia National Laboratories, a multimission laboratory managed and operated by National Technology and Engineering Solutions of Sandia LLC, a wholly owned subsidiary of Honeywell International Inc. for the U.S. Department of Energy’s National Nuclear Security Administration under contract DE-NA0003525.

## Appendix

### A Supplementary Materials

For further experimentation, it is important to recognize that scientists and researchers rarely interact with LLMs through single, isolated prompts. Instead, they typically engage in multi-turn conversations, iteratively refining their queries to guide the model toward the desired output. In this study, our initial evaluations used single-shot prompts to assess model responses. To better simulate real-world usage, we conducted an additional experiment with two models, Claude Sonnet 4.6 and ChatGPT 5.5, in which we used the most specific (Version 3) prompts for the Full Test and LOF tasks and allowed up to ten conversational turns. This iterative approach aimed to bring the LLM-generated outputs and plots closer to those produced by the hard-coded pipeline.

The results of this conversational experiment are presented below.

#### A.1 Pipeline Gold Standards

##### Experiment 4. Full Test

Pipeline Silhouette: 0.5582

Pipeline ARI: 0.4129

##### Experiment 5. LOF

Pipeline outlier count: 58

A.2 ChatGPT 5.5

### Full Test

As can be seen in Table 16, in this iterative refinement experiment, the model required only two additional conversational turns to produce outputs that more closely matched the reference pipeline results.

**Table 15:** Reproducibility results for Llama4-Scout across prompt versions.

| Version | Run 1 Silhouette | Run 2 Silhouette | Match? |
| --- | --- | --- | --- |
| v1 | 0.8584 | 0.8584 | Yes |
| v2 | 0.8584 | 0.8584 | Yes |
| v3 | 0.8584 | 0.8584 | Yes |

**Table 16:** Iterative refinement results for ChatGPT 5.5 on the Full Test task.

| Turn # | Input Prompt | Silhouette Score | ARI | Notes |
| --- | --- | --- | --- | --- |
| 1 | Based on the attached file, the corresponding silhouette score and ari produced were very different from the pipeline scores (silhouette: 0.5582 and ari: 0.4129) and the produced plot contained 4 clusters rather than three, why is this the case and can you tweak the script to potentially fix it | 0.463 | 0.249 | ARI score is significantly closer, silhouette is slightly closer to pipeline but still not exact. No difference in output plot. |
| 2 | The scores are closer, but still not exact. The plot is still off, with four clusters rather than the three. Please give me ideas on what is causing the discrepancy and proposed modified script. | 0.558 | 0.413 | Nearly identical scores, both silhouette and ARI, and nearly identical plot |

During this process, the model was able to identify and correct several discrepancies between its initial standalone script and the implemented pipeline. The revised script was modified to use the pipeline-ready sample-by-gene CSV input format, apply the same column-normalization procedure used by CSV_Pandas, and align PCA preprocessing with the pipeline by selecting only numeric features and imputing missing values with feature means. The dimensionality-reduction and clustering sequence was also revised to match the pipeline implementation: PCA was performed first, t-SNE was applied to the PCA-transformed data, and KMeans clustering was then performed on the resulting t-SNE coordinates.

These revisions were important because the model’s initial script performed clustering in PCA space while visualizing the results in t-SNE space, whereas the reference pipeline clustered directly on the t-SNE embedding. Aligning the preprocessing steps, clustering space, and order of operations reduced discrepancies between the standalone script and the pipeline-derived outputs, including the Silhouette Score, Adjusted Rand Index (ARI), and visual cluster structure. Remaining differences may be attributable to stochastic t-SNE initialization, default parameter differences across scikit-learn versions, and potential study or batch effects in the combined four-study dataset.

### LOF

It can be seen in Table 17 that the model again required only one additional conversational turn to generate a script that more closely matched the reference pipeline output. The revised script recognized that reproducing the pipeline’s outlier-detection results required matching the pipeline’s intermediate data flow rather than independently reconstructing all preprocessing steps from the raw input data. Specifically, the updated script first attempts to load the actual PCA output pickle generated by the pipeline from comms_files and then applies the same LOFAndFiltering logic to that PCA matrix.

**Table 17:** Iterative refinement results for ChatGPT 5.5 on the LOF task.

| Turn # | Input Prompt | Outlier Count | Notes |
| --- | --- | --- | --- |
| 1 | Based on the attached file, the corresponding outlier count of 58 was slightly different from the pipeline outline count of 56. Explain why this may be the case and can you tweak the script to potentially fix it. | 58 | Correct outlier count! |

This change was important because the original difference in outlier count was likely caused by small numerical or preprocessing differences between the standalone script and the pipeline. Potential sources of discrepancy include mean-imputation behavior, feature ordering, PCA implementation details, random-state handling, or borderline LOF scores near the cutoff value of 1.5. To support debugging and reproducibility, the revised script also added diagnostic reporting for strict cutoff-based outliers, LOF model-predicted outliers, and samples with scores closest to the threshold. These diagnostics make it easier to determine whether any remaining discrepancy is caused by rounding, threshold sensitivity, or near-boundary outlier scores.

#### A.3 Claude Sonnet 4.6

##### Full Test

In this additional iterative-refinement experiment for the Full Test task outlined in Table 18, this model performed substantially worse than ChatGPT 5.5. Despite receiving corrective feedback across the allowed interaction window, it was unable to produce outputs that matched the reference pipeline results within the 10-turn limit.

**Table 18:** Iterative refinement results for Claude Sonnet 4.6 on the Full Test task.

| Turn # | Input Prompt | Silhouette Score | ARI | Notes |
| --- | --- | --- | --- | --- |
| 1 | Based on the attached file, the corresponding silhouette score and ari produced were very different from the pipeline scores (silhouette: 0.5582 and ari: 0.4129) and the produced plot contained 4 clusters rather than three, why is this the case and can you tweak the script to potentially fix it |  |  | Code errored out. |
| 2 | max_iter should be replaced by n_iter | 0.858 | -0.00700 | Scores didn't change, still very different from pipeline. |
| 3 | Once again, the silhouette score produced was 0.8584 and ari was -0.0070 with 4 clustered plot rather than 3 (pipeline produced plot) and the pipeline silhouette score was 0.5582 and Pipeline ARI: 0.4129. I want the script you produce to be modified to be closer to this, explain why there would be a discrepancy and then provide the modifications as a whole new script | 0.452 | 0.199 | Closer scores to pipeline but still not identical. Plot is still not the same as pipeline plot. |
| 4 | the metrics this script produced are as follows: 0.4517 ARI vs true class_label: 0.1933 and the plot still shows 4 clusters rather than 3, please provide a further explanation and fix | 0.858 | -0.00700 | Back to original incorrect scores, plot also incorrectly clustered into 4 rather than 3. |
| 5 | Attached is the pipeline produced full test-plot. As a reminder the pipeline silhouette score was 0.5582 and ARI 0.4129. This latest script produced the original incorrect silhouette score of 0.8584 and ARI of -0.0070. What is the reason for this and please fix this. | 0.222 | -0.00700 | Same incorrect ARI, different silhouette score, still incorrect. Same incorrect plot. |
| 6 | Not quite. I will attach an image of the plot that this latest script produced. Completely different, fix this. Additionally, this did not change the ARI, only the silhouette score to 0.2217 which is still incorrect. | 0.460 | 0.245 | Much improved scores and closer plot, still not identical to pipeline. |
| 7 | Almost. The scores produced were as followed: Silhouette Score : 0.4604 ARI : 0.2487 with the attached plot. Is there anyway to get it closer to the pipeline results? | 0.458 | 0.187 | Incorrect scores and plot. |
| 8 | This is the input csv file |  |  | Took too long to load response; no script produced. |
| 9 | This was the latest silhouette and ARI: Silhouette Score : 0.4584 (pipeline: 0.5582) ARI : 0.1871 (pipeline: 0.4129) with the same plot as from the previous plot. Another fix? |  |  | Took too long to load response; no script produced. |
| 10 | It's taking a long time, ignore the csv and just pay attention to what i said: This was the latest silhouette and ARI: Silhouette Score : 0.4584 (pipeline: 0.5582) ARI : 0.1871 (pipeline: 0.4129) with the same plot as from the previous plot. Another fix? | 0.474 | 0.0616 | Very different scores, different but still incorrect plot. |

##### LOF

In the LOF experiment outlined in Table 19, the model required three additional conversational turns to adjust its output so that it more closely matched the reference pipeline results. During refinement, the model identified that the main discrepancy came from differences between the standalone implementation and the pipeline’s chained component structure. In the reference workflow, PCAnalysis first generates the PCA representation, and LOFAndFiltering then operates on that PCA output rather than independently preprocessing and analyzing the original feature matrix.

**Table 19:** Iterative refinement results for Claude Sonnet 4.6 on the LOF task.

| Turn # | Input Prompt | Outlier Count | Notes |
| --- | --- | --- | --- |
| 1 | Based on the attached file, the corresponding outlier count of 58 was slightly different from the pipeline outline count of 56. Explain why this may be the case and can you tweak the script to potentially fix it. I will also attach the pipeline generated plot for LOF as further guidance. | 94 | Very different outlier count, farther away than the original v3 script produced. |
| 2 | Not quite, this was the output produced, it was even farther away from the pipeline produced number than the first attached .py file. I will attach the component_library file that is used in the pipeline- I used the LOFAndFiltering component. | 118 | Farther away from correct outlier count than previously |
| 3 | This produced an outlier count of 118, worse than before | 56 | Slightly outlier count |

To better align with the pipeline, the revised script incorporated three key changes. First, it removed the StandardScaler step because the pipeline’s PCAnalysis component passes the raw gene-expression values directly into PCA without applying explicit normalization. Second, it removed the fixed random_state parameter from PCA because the pipeline’s PCA() call does not specify a seed; setting random_state=42 in the standalone script produced a different component space and, consequently, different LOF score distributions. Third, the script was modified so that LOF was applied to the PCA output DataFrame rather than to the original high-dimensional gene-expression matrix. In the pipeline, PCAnalysis returns a principalDf containing 50 principal-component columns, along with a label column that is subsequently excluded by numeric-column selection, and this 50-dimensional PCA representation is the input used by LOFAndFiltering.

The missing-value handling was also revised to match the pipeline more closely. Instead of using fillna(0), the updated script drops columns containing only missing values and then imputes remaining missing entries using feature means, consistent with the preprocessing performed by PCAnalysis before transformation. Together, these changes reduced discrepancies between the standalone script and the pipeline by aligning the preprocessing assumptions, PCA configuration, intermediate data representation, and LOF input space.

## Supplementary Experiment Conclusions

Overall, within the 10-turn refinement limit, ChatGPT 5.5 produced outputs that aligned more closely with the reference pipeline than Claude Sonnet 4.6 and required fewer conversational turns to do so. This suggests that, in this small-sample experiment, ChatGPT 5.5 was more effective at using corrective feedback to identify discrepancies between its standalone scripts and the expected pipeline behavior. In particular, it more readily adjusted preprocessing choices, intermediate data flow, and analysis ordering to match the implemented biological data-analysis workflow.

However, because the experiment used a limited number of tasks and a small sample of model interactions, the finding should not be taken as a general ranking of model capability. Rather, it indicates that, for the specific pipeline-reproduction tasks tested here, ChatGPT 5.5 demonstrated stronger iterative alignment with the reference workflow than Claude Sonnet 4.6 under the same 10-turn constraint. This supports the broader conclusion that conversational refinement can improve LLM-generated scientific workflows, but that the degree of improvement is model-dependent and still relies heavily on knowledgeable user feedback.

